# Development of single-molecule enzyme activity assay for serine hydrolases using activity-based protein labeling probes

**DOI:** 10.1101/2025.08.06.668846

**Authors:** Seiya Ishii, Mayano Minoda, Tadahaya Mizuno, Takumi Iwasaka, Hiroyuki Kusuhara, Kazufumi Honda, Yasuteru Urano, Toru Komatsu

## Abstract

Single-molecule enzyme activity assays have proven their potential in elucidating aberrant protein function at the proteoform level. However, the limited number of targetable enzymes is the major drawback of such assays. Here, we report the development of single-molecule enzyme activity assays utilizing activity-based probes that label active enzymes in an enzyme superfamily-wide manner. A proof-of-principle using fluorophosphonate-based probes was conducted to detect the active form of serine hydrolases such as PSA and granzyme B at the single-molecule level in complex biological systems such as blood. The assay suggested the potential of active granzyme B in blood as a suitable biomarker of liver damage with specific activation of immune cells.

## MAIN TEXT

In living systems, there are thousands of enzymes involved in maintaining homeostasis, and their alterations are often tightly connected to the onset of diseases^1–4^. Enzyme activities are controlled at the transcriptional, translational, and posttranslational levels, and analysis of their functional alterations is useful for detecting and understanding diseases^3,4^. Single-molecule enzyme activity analysis of protein function shows high sensitivity, and has proven useful in detecting disease-related alteration of functional states of proteins at the proteoform level^5–9^. The present single-molecule enzyme assay employs fluorogenic substrate probes that are metabolized by enzymes to become fluorescent. However, the class of enzymes to which fluorogenic substrates produce sufficient signals remains limited^9^, and the requirement for individual development of specific substrates is the bottleneck to broad assay deployment. In this study, we propose a general methodology that utilizes activity-based protein labeling probes (ABPs) to detect various single-molecule enzyme activities in an enzyme superfamily-selective manner, by selectively labeling enzymes with accessible active sites (**Figure 1**).

**Figure 1.**
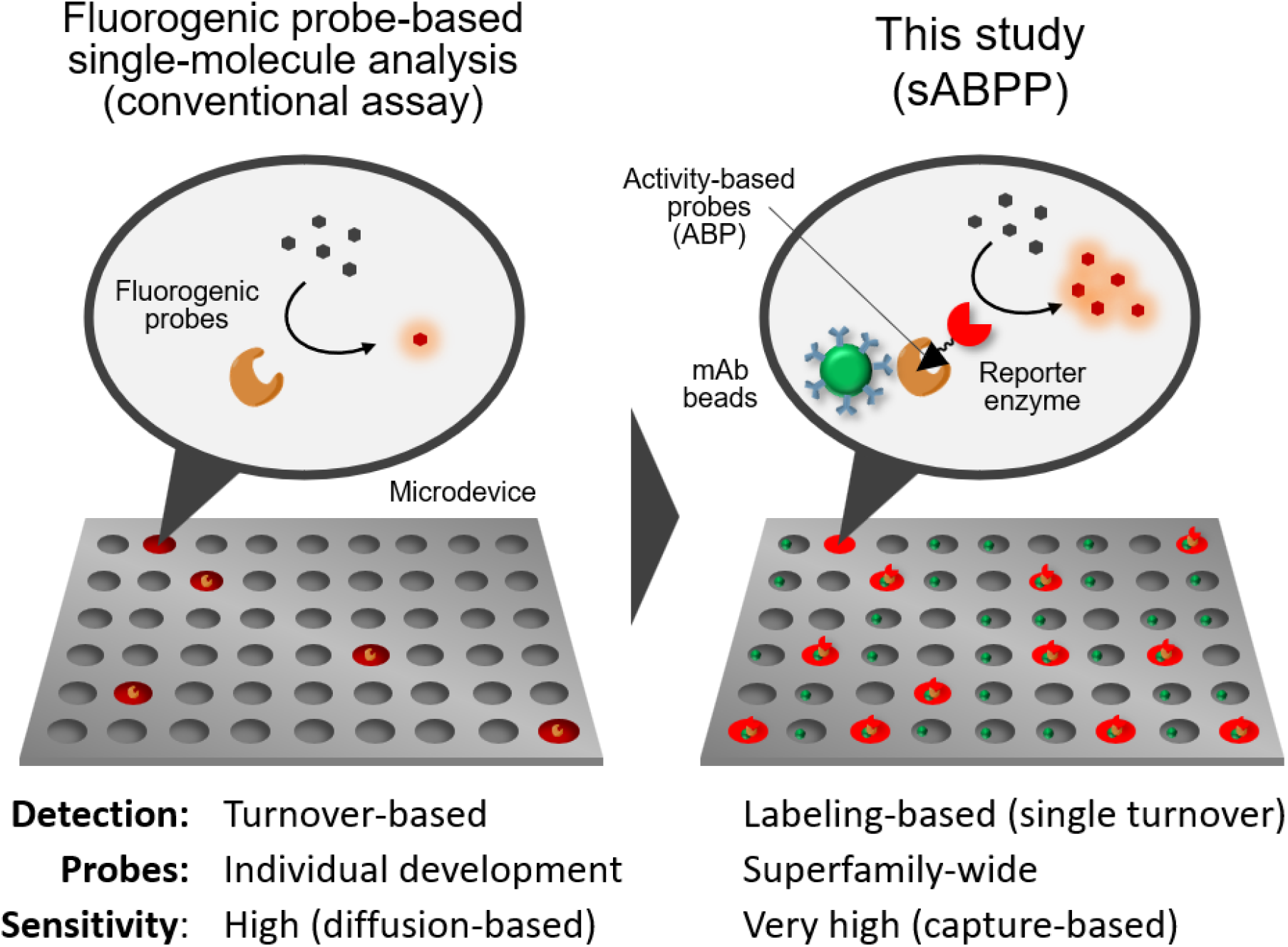
Schematic view of single-molecule enzyme activity-based protein profiling (sABPP) and its advantages compared to conventional fluorogenic probe-based single-molecule enzyme activity assays.

ABPs are molecules that covalently label the active site of enzymes, enabling discrimination between active and inactive enzymes. Activity-based protein profiling (ABPP) utilizes these characteristics for global analysis of enzyme activities^10,11^, and we propose that single-molecule enzyme activity assays can be performed by detecting ABP-based labeling at the single-molecule level, which enables selective detection of enzymes with accessible active sites. The single-molecule enzyme activity assay is performed by capturing the single-molecule target enzyme in a microfabricated chamber device, and the metabolism of the fluorogenic substrate generates a fluorescent signal in the chamber. The detection relies on enzymatic amplification of the signal, such that covalently attaching a reporter enzyme to the ABP can realize its readout. Since ABPs globally label various enzymes in the same superfamily, we combined the ABP with antibody-based capture of the target enzyme for selective detection of the protein of interest (**Figure S1**).

We initiated the proof-of-concept study of single-molecule activity-based protein profiling (sABPP) using prostate-specific antigen (PSA) as a biomarker of prostate cancer^12^.

Currently, antibody-based detection is commonly used to measure PSA in blood as a biomarker of prostate cancer, and single-molecule assays such as digital ELISA have also been applied; however, these methods detect the total amount of PSA molecules regardless of their enzymatic activity, and a lack of attention has been paid to alterations in its activity. PSA (kallikrein 3) is a chymotrypsin-like endopeptidase, and its activity contributes to various physiological processes such as reproduction, regulation of blood pressure, and inflammation^13^. The enzyme is generated as a preproenzyme and must be correctly processed by cell surface proteases to become proteolytically active^14^.

Serine hydrolases can be labeled at their active sites by fluorophosphonate (FP)^10,15^. Using FP-based ABPs, we constructed a system to selectively detect the “active” form of PSA at the single-molecule level (**Figure 2a**). After the activity-based labeling of PSA with FP-biotin, it was captured by an anti-PSA antibody bound to fluorescent magnetic beads (green fluorescence; labeled by Alexa Fluor 488), and biotin was labelled with streptavidin β-Gal (SA β-Gal). The complex was loaded into a microdevice and the number of chambers containing β-galactosidase activity was read out by the fluorescence of resorufin β-gal (red fluorescence). Loading of the complex was automated using the digital ELISA instrument siMoA^16^, and detection of the fluorescent complexes was performed using a standard epifluorescence microscope. The assay selectively detected PSA activated by thermolysin (active-form, **Figure S2**), as the increase of red fluorescent spots reflecting β-galactosidase activity. The spot number was significantly lower for the preproenzymes (pro-form) or enzymes treated with inhibitors (**Figure 2b, 2c**).

**Figure 2.**
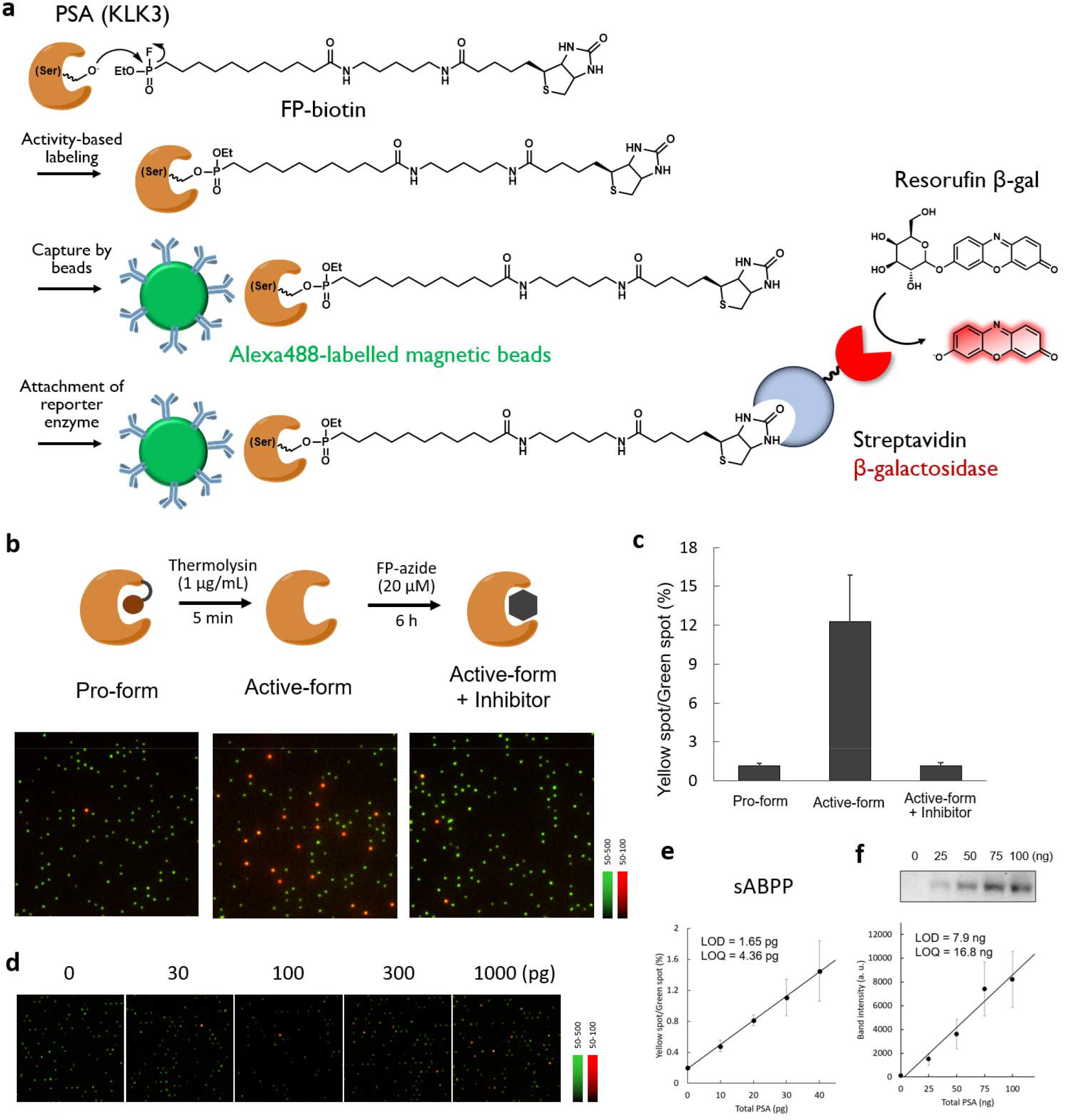
Construction of single-molecule enzyme activity-based protein profiling assay using PSA. (a) PSA labeling scheme for activity-based probes, antibody capture, and streptavidin β-Gal attachment for activity readout. (b) Detection of single-molecule active PSA in the pro-enzyme form, active-form, and inhibitor-treated active-form samples. (c) Quantification of the result of **Figure 2b**. Yellow spots refer to the particles comprised of green fluorescent beads that produced red fluorescence following resorufin production. Green spots refer to the particles comprised of green fluorescent beads with no resorufin production. (d) Fluorescence images of the microdevice loaded with various amounts of activated PSA. The labeling and loading conditions were the same as those in **Figure 2b**. (e) Correlation between the proportion of yellow spots and activated PSA in **Figure 2d**. The amount of PSA refers to pro-PSA used for the assay. Error bars represent S. D. (n = 3). Limit of detection (LOD) was calculated as the value corresponding to 3σ. Limit of quantification (LOQ) was calculated as the value corresponding to 10σ. (f) LOD and LOQ calculated in ABP labeling and chemiluminescence-based detection by western blotting. Error bars represent S. D. (n = 3). The same protein samples were used in **Figure 2d** and **2e**. The amount of PSA refers to pro-PSA used for activation.

The advantage of the single-molecule assay is its high detection sensitivity compared with conventional protein detection platforms performed under bulk conditions. We confirmed that the assay has a limit of detection (LOD) that is >1,000-times lower than SDS-PAGE/chemiluminescence detection of the same protein sample (**Figure 2d-2f**). The high sensitivity is attributed to the combination of the high sensitivity of the single-molecule analysis itself and antibody enrichment of the target (**Figure 1**). It should be noted that, in this assay, PSA is detected after activation from its proenzyme form, and since the activation efficiency is undefinable, comparison with the LOD of commercial total PSA assays is not straightforward. Nonetheless, the detection sensitivity for active PSA is expected to be comparable to that of the digital ELISA assay for total PSA^17^, since the enrichment by antibody beads operates under the same principle. The assay showed sufficient selectivity toward other serine hydrolases, such as trypsin and elastase, based on the selectivity of the capture antibody (**Figure S1**); proteases that were labeled by FP-biotin, such as trypsin and elastase, were not captured by the antibody-modified beads and did not generate red fluorescent spots (**Figure 3a-3c**).

**Figure 3.**
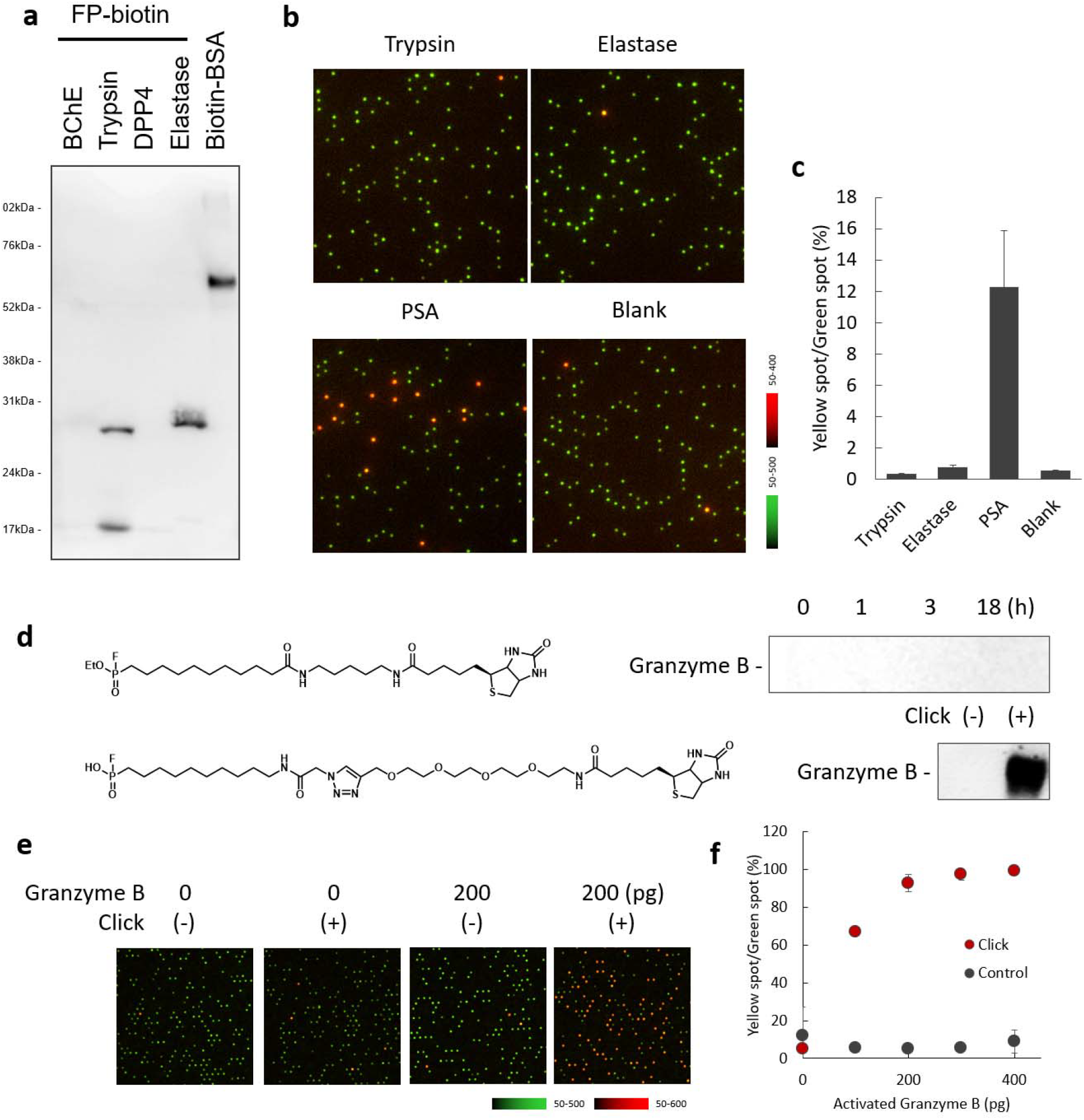
Applicability of sABPP to other serine hydrolases. (a) Western blot analysis of serine hydrolases labeled with FP-biotin. Biotin-modified BSA was used as positive control. (b) Representative fluorescence overlay images of the microdevice containing recombinant enzymes (trypsin, elastase, activated PSA) or blank. The protocol was the same as that of **Figure 2b**. (c) Quantification of the result of **Figure 3b**. (d) FP-biotin probes with different linker lengths for granzyme B labeling. Western blot analysis of activated granzyme B. For labeling with FP-biotin (top), the labeling/detection condition was the same as that in **Figure 3a**. For labeling with FP-azide (bottom), granzyme B was labeled by FP-azide, and the sample was reacted with alkyne-PEG4-biotin for click-reaction. (e) Detection of recombinant granzyme B using FP-azide. The labeling was performed under the same condition as in **Figure 3d**. (f) Quantification of the result in Figure 3e. The amount of activated granzyme B refers to the granzyme B used for the assay. Error bars represent S. D. (n = 3).

Due to the generality of the assay scheme, it can target various serine hydrolases by changing the capture antibodies. We next developed another single-molecule activity assay targeting granzyme B, which is generated by natural killer (NK) cells and CD8^+^ T cells^18^. Due to its weak reactivity and scarcity in blood, there are no reports of granzyme B activity detection in blood samples. As granzyme B is also a serine hydrolase, we predicted that active granzyme B could be detected by FP-based labeling. The detection of granzyme B required the optimization of the ABP linker length; ABP with a short linker (FP-biotin) failed to form a complex with granzyme B and streptavidin, so we extended the linker between FP and biotin after labeling via click chemistry (FP-azide + biotin-PEG4-alkyne), which successfully enabled complex formation with streptavidin (**Figure 3d**). We consider that this is due to structural constraints, such as the depth of the granzyme B binding pocket and steric hindrance around streptavidin, preventing the two proteins from coming close enough when using ABPs with short linkers^19^. By employing the click-chemistry-based linker extension strategy, we were able to detect active granzyme B at the single-molecule level (**Figure 3e, 3f**).

We then determined if active PSA and granzyme B are present in blood in an effort to develop them as activity-based biomarkers. Active PSA was not observed in plasma samples from healthy human subjects (**Figure 4a**). When total PSA in the same blood sample was measured using a detection antibody conjugated with the reporter enzyme (**Figure 4b**)^17^, red fluorescent spots reflecting the secondary antibody were observed. The result indicates that “inactive” PSA was the dominant form in blood due to the lack of activation by processing enzymes or inhibition by anti-proteases (e.g. serpins)^14^. Active PSA was spiked into plasma samples before labeling, and the number of red fluorescent spots decreased, supporting the idea that anti-proteases for PSA are dominant in blood samples of healthy human subjects (**Figure S3**). In contrast, active granzyme B was detected in plasma samples of healthy human subjects (**Figure 4c**). We determined whether granzyme B activity in blood samples is altered under different health conditions, with the aim of developing an activity-based disease biomarker. Since granzyme B is generated exclusively in natural killer (NK) cells and CD8^+^ T cells, we expected that its activity can be altered by specific immune responses. To investigate this idea, we prepared two mouse models of liver damage by treating subjects with thioacetamide (TAA)^8,20^ or 4,4’-methylenedianiline (MDA)^21^, which are known to cause different forms of immune responses in liver. The TAA-model leads to acute liver damage and gradual liver fibrosis^20^, while the MDA-model leads to increased monocyte-derived macrophages and enhanced fibrinolysis in liver^21,22^. While the mechanisms are different, the present biomarker AST/ALT, which is elevated during liver damage as tissue leakage products, cannot discriminate between these two liver damage models (**Figure S4**)^22^. Interestingly, elevated levels of active granzyme B were observed exclusively in blood samples of MDA-treated mice (**Figure 4d, 4e, S5**). We interpreted this as a reflection of an increase in CD8^+^ T cells in MDA-treated mice^22^. The overall results indicate that activity-based biomarker detection by single-molecule enzyme activity assay can selectively detect specific forms of liver damage. Since granzyme B activity is important for tumor immunity^23^, we contend that alterations in granzyme B activity can serve as a meaningful blood biomarker to evaluate changes in immune status of tumor tissues, thereby contributing to early tumor detection and better classification of patients that respond to immune checkpoint inhibitors.

**Figure 4.**
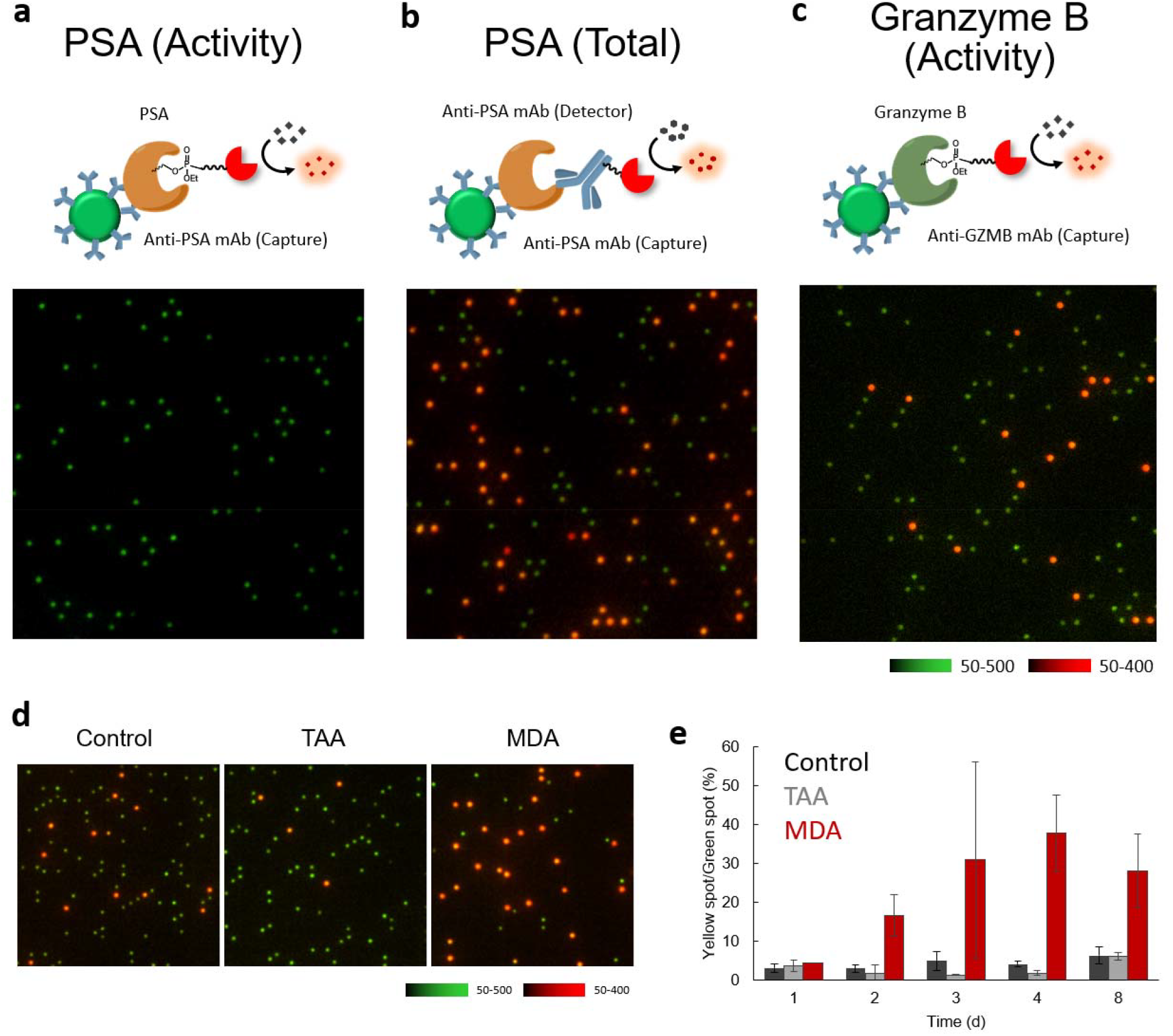
Analysis of serine hydrolase activities in blood using sABPP. (a) Detection of active PSA in plasma samples from healthy human subjects (male, 1/300 dilution) using the same condition as in **Figure 2b**. (b) Detection of total PSA in plasma samples from healthy human subjects (male, 1/300 dilution) using a digital ELISA platform. Biotin-labeled secondary antibody was used instead of labeling with FP-biotin. (c) Detection of active granzyme B in plasma samples from healthy human subjects (male, 1/300 dilution) using the same condition as in **Figure 3e**. (d) Detection of active granzyme B in plasma samples of mice treated with/without thioacetamide (TAA) as an acute liver injury model or 4,4’-methylene dianiline (MDA) as a cholestatic liver dysfunction model. Plasma samples collected at 4 days of treatment were used. The plasma samples were diluted 1/300 for the assay. (e) Time-course changes of active granzyme B in liver damage models. Error bars represent S. D. (n = 3).

In summary, we report a single-molecule activity-based protein profiling (sABPP) platform that utilizes ABPs to selectively detect active enzymes at the single-molecule level. A wide variety of ABPs are currently being developed for enzyme families such as serine hydrolases, cysteine hydrolases, glycosidases, phosphatases, and kinases^10,11^, and their use in mapping the reactive landscape of proteins has also been proposed^24–26^. In combination with these advances in ABP development, the assay scheme presented here has the potential to expand the range of protein activities accessible to single-molecule analysis. In the future, multiplexed detection of related enzymatic activities and reactivities at the single-molecule level will likely become feasible, increasing the likelihood of identifying disease-associated proteoforms. Overall, the system provides a sensitive and versatile framework for detecting functional proteins relevant to disease diagnosis and the elucidation of biological mechanisms.

## ASSOCIATED CONTENT

### Supplementary Information

Methods and supporting tables/figures.

## AUTHOR INFORMATION

### Author Contribution

T. K. conceived the experimental design. S. I. and M. M. acquired experimental data. T. M., T. I., and H. K. prepared plasma samples from mice. K.H. prepared plasma samples from human healthy subjects. The experimental data were analyzed under the supervision of Y. U. and T. K. The manuscript was written by T. K.

## ACKNOWLEDGMENT

We thank Dr. Ayumu Kashiro (Nippon Medical School) and Dr. Oketani (Kagoshima Prefectural Comprehensive Health Center) for collecting the plasma samples.

## REFERENCES

(1) Komatsu, T.; Urano, Y. Evaluation of Enzymatic Activities in Living Systems with Smallmolecular Fluorescent Substrate Probes. Anal. Sci. 2015, 31 (4), 257–265. 10.2116/analsci.31.257.

(2) Komatsu, T.; Urano, Y. Chemical Toolbox for “live” Biochemistry to Understand Enzymatic Functions in Living Systems. J. Biochem. Oxford University Press August 15, 2020, pp 139–149. 10.1093/jb/mvz074.

(3) Soleimany, A. P.; Bhatia, S. N. Activity-Based Diagnostics: An Emerging Paradigm for Disease Detection and Monitoring. Trends Mol. Med. 2020, 26 (5), 450–468. 10.1016/j.molmed.2020.01.013.

(4) Saghatelian, A.; Cravatt, B. F. Assignment of Protein Function in the Postgenomic Era. Nat. Chem. Biol. 2005, 1 (3), 129. 10.1038/nchembio0805-130.

(5) Nagano, N.; Ichihashi, Y.; Komatsu, T.; Matsuzaki, H.; Hata, K.; Watanabe, T.; Misawa, Y.; Suzuki, M.; Sakamoto, S.; Kagami, Y.; Kashiro, A.; Takeuchi, K.; Kanemitsu, Y.; Ochiai, H.; Watanabe, R.; Honda, K.; Urano, Y. Development of Fluorogenic Substrates for Colorectal Tumor-Related Neuropeptidases for Activity-Based Diagnosis. Chem. Sci. 2023. 10.1039/d2sc07029d.

(6) Sakamoto, S.; Hiraide, H.; Minoda, M.; Iwakura, N.; Suzuki, M.; Ando, J.; Takahashi, C.; Takahashi, I.; Murai, K.; Kagami, Y.; Mizuno, T.; Koike, T.; Nara, S.; Morizane, C.; Hijioka, S.; Kashiro, A.; Honda, K.; Watanabe, R.; Urano, Y.; Komatsu, T. Identification of Activity-Based Biomarkers for Early-Stage Pancreatic Tumors in Blood Using Single-Molecule Enzyme Activity Screening. Cell Rep. Methods 2024, 4, 100688. 10.1016/j.crmeth.2023.100688.

(7) Sakamoto, S.; Komatsu, T.; Watanabe, R.; Zhang, Y.; Inoue, T.; Kawaguchi, M.; Nakagawa, H.; Ueno, T.; Okusaka, T.; Honda, K.; Noji, H.; Urano, Y. Multiplexed Single-Molecule Enzyme Activity Analysis for Counting Disease-Related Proteins in Biological Samples. Sci. Adv. 2020, 6 (11), eaay0888. 10.1126/sciadv.aay0888.

(8) Ukegawa, T.; Komatsu, T.; Minoda, M.; Matsumoto, T.; Iwasaka, T.; Mizuno, T.; Tachibana, R.; Sakamoto, S.; Hanaoka, K.; Kusuhara, H.; Honda, K.; Watanabe, R.; Urano, Y. Thioester-Based Coupled Fluorogenic Assays in Microdevice for the Detection of Single-Molecule Enzyme Activities of Esterases with Specified Substrate Recognition. Adv. Sci. 2023, 11 (10), 2306559. 10.1002/advs.202306559.

(9) Komatsu, T.; Mizuno, T. Single-Molecule Enzyme Activity Analysis for Illuminating Pathological Proteoforms. ACS Cent. Sci. 2025, 11 (7), 1041–1051. 10.1021/acscentsci.5c00100.

(10) Liu, Y.; Patricelli, M. P.; Cravatt, B. F. Activity-Based Protein Profiling: The Serine Hydrolases. Proc. Natl. Acad. Sci. USA 1999, 96 (26), 14695–14699.

(11) Nomura, D. K.; Dix, M. M.; Cravatt, B. F. Activity-Based Protein Profiling for Biochemical Pathway Discovery in Cancer. Nat. Rev. Cancer 2010, 10 (9), 630–638. 10.1038/nrc2901.

(12) Papsidero, L. D.; Wang, M. C.; Valenzuela, L. A.; Murphy, G. P.; Chu, T. M. A Prostate Antigen in Sera of Prostatic Cancer Patients1. Cancer Res. 1980, 40, 2428–2432.

(13) Lawrence, M. G.; Lai, J.; Clements, J. A. Kallikreins on Steroids: Structure, Function, and Hormonal Regulation of Prostate-Specific Antigen and the Extended Kallikrein Locus. Endocr. Rev. 2010, 31 (4), 407–446. 10.1210/er.2009-0034.

(14) Hori, S.; Blanchet, J. S.; McLoughlin, J. From Prostate-Specific Antigen (PSA) to Precursor PSA (ProPSA) Isoforms: A Review of the Emerging Role of ProPSAs in the Detection and Management of Early Prostate Cancer. BJU Int. 2013, 112 (6), 717–728. 10.1111/j.1464-410X.2012.11329.x.

(15) Bachovchin, D. A.; Ji, T.; Li, W.; Simon, G. M.; Blankman, J. L.; Adibekian, A.; Hoover, H.; Niessen, S.; Cravatt, B. F. Superfamily-Wide Portrait of Serine Hydrolase Inhibition Achieved by Library-versus-Library Screening. Proc. Natl. Acad. Sci. USA 2010, 107 (49), 20941–20946. 10.1073/pnas.1011663107/-/DCSupplemental.

(16) Kan, C. W.; Rivnak, A. J.; Campbell, T. G.; Piech, T.; Rissin, D. M.; Mösl, M.; Petera, A.; Niederberger, H. P.; Minnehan, K. A.; Patel, P. P.; Ferrell, E. P.; Meyer, R. E.; Chang, L.; Wilson, D. H.; Fournier, D. R.; Duffy, D. C. Isolation and Detection of Single Molecules on Paramagnetic Beads Using Sequential Fluid Flows in Microfabricated Polymer Array Assemblies. Lab Chip 2012, 12 (5), 977–985. 10.1039/c2lc20744c.

(17) Rissin, D. M.; Kan, C. W.; Campbell, T. G.; Howes, S. C.; Fournier, D. R.; Song, L.; Piech, T.; Patel, P. P.; Chang, L.; Rivnak, A. J.; Ferrell, E. P.; Randall, J. D.; Provuncher, G. K.; Walt, D. R.; Duffy, D. C. Single-Molecule Enzyme-Linked Immunosorbent Assay Detects Serum Proteins at Subfemtomolar Concentrations. Nat. Biotechnol. 2010, 28 (6), 595–599. 10.1038/nbt.1641.

(18) Scott, J. I.; Mendive-Tapia, L.; Gordon, D.; Barth, N. D.; Thompson, E. J.; Cheng, Z.; Taggart, D.; Kitamura, T.; Bravo-Blas, A.; Roberts, E. W.; Juarez-Jimenez, J.; Michel, J.; Piet, B.; de Vries, I. J.; Verdoes, M.; Dawson, J.; Carragher, N. O.; Connor, R. A. O.; Akram, A. R.; Frame, M.; Serrels, A.; Vendrell, M. A Fluorogenic Probe for Granzyme B Enables In-Biopsy Evaluation and Screening of Response to Anticancer Immunotherapies. Nat. Commun. 2022, 13 (1). 10.1038/s41467-022-29691-w.

(19) Sato, S. I.; Kwon, Y.; Kamisuki, S.; Srivastava, N.; Mao, Q.; Kawazoe, Y.; Uesugi, M. Polyproline-Rod Approach to Isolating Protein Targets of Bioactive Small Molecules: Isolation of a New Target of Indomethacin. J. Am. Chem. Soc. 2007, 129 (4), 873–880. 10.1021/ja0655643.

(20) Wallace, M.; Hamesch, K.; Lunova, M.; Kim, Y.; Weiskirchen, R.; Strnad, P.; Friedman, S. L. Standard Operating Procedures in Experimental Liver Research: Thioacetamide Model in Mice and Rats. Lab. Anim. 2015, 49, 21–29. 10.1177/0023677215573040.

(21) Kwon, S. B.; Park, J. S.; Yi, J. Y.; Hwang, J. W.; Kim, M.; Lee, M. O.; Lee, B. H.; Kim, H. L.; Kim, J. H.; Chung, H.; Kong, G.; Kang, K. S.; Yoon, B. Il. Time- and Dose-Based Gene Expression Profiles Produced by a Bile-Duct-Damaging Chemical, 4,4′-Methylene Dianiline, in Mouse Liver in an Acute Phase. Toxicol. Pathol. 2008, 36 (5), 660–673. 10.1177/0192623308320272.

(22) Iwasaka, T.; Mizuno, T.; Morita, K.; Azuma, I.; Nakagawa, T.; Nakashima, E.; Kusuhara, H. Establishment of an Easy-to-Construct Liver Injury Mouse Model for Longitudinal Analysis by Drinking-Water Administration of MDA. bioRxiv 2024. 10.1101/2024.01.25.577198.

(23) Sharma, P.; Allison, J. P. The Future of Immune Checkpoint Therapy. Science (1979) 2015, 348 (6230), 56–61.

(24) Weerapana, E.; Wang, C.; Simon, G. M.; Richter, F.; Khare, S.; Dillon, M. B. D.; Bachovchin, D. A.; Mowen, K.; Baker, D.; Cravatt, B. F. Quantitative Reactivity Profiling Predicts Functional Cysteines in Proteomes. Nature 2010, 468 (7325), 790–797. 10.1038/nature09472.

(25) Bar-Peled, L.; Kemper, E. K.; Suciu, R. M.; Vinogradova, E. V.; Backus, K. M.; Horning, B. D.; Paul, T. A.; Ichu, T. A.; Svensson, R. U.; Olucha, J.; Chang, M. W.; Kok, B. P.; Zhu, Z.; Ihle, N. T.; Dix, M. M.; Jiang, P.; Hayward, M. M.; Saez, E.; Shaw, R. J.; Cravatt, B. F. Chemical Proteomics Identifies Druggable Vulnerabilities in a Genetically Defined Cancer. Cell 2017, 171 (3), 696–709.e23. 10.1016/j.cell.2017.08.051.

(26) Abbasov, M. E.; Kavanagh, M. E.; Ichu, T. A.; Lazear, M. R.; Tao, Y.; Crowley, V. M.; am Ende, C. W.; Hacker, S. M.; Ho, J.; Dix, M. M.; Suciu, R.; Hayward, M. M.; Kiessling, L. L.; Cravatt, B. F. A Proteome-Wide Atlas of Lysine-Reactive Chemistry. Nat. Chem. 2021, 13 (11), 1081–1092. 10.1038/s41557-021-00765-4.

