## Supplementary_Information for "Development of single-molecule enzyme activity assay for serine hydrolases using activity-based protein labeling probes"

**Contents**

**Methods**
**Supplementary figures**

### Methods

#### Plasma samples from healthy human subjects and ethics statement

Plasma samples from healthy human subjects were collected from Kagoshima Prefectural Comprehensive Health Center with the Program for Promotion of Fundamental Studies in Health Sciences conducted by the National Institute of Biomedical Innovation of Japan, Health and Labour Sciences Research Grants from the Ministry of Health, Labour and Welfare of Japan, and P-CREATE of the Japan Agency for Medical Research and Development (AMED). Ethical approval for this study was obtained from the central ethics committees of Nippon Medical School (M-2021-002) and the ethical committee of Nippon Medical School (A-2020-032 and A-2020-044).

#### Plasma samples from mice and ethics statement

Ethical approval for the study using animals was obtained from the Animal Care and Use Committee of The University of Tokyo (P4-21, P31-9). Six-week-old male C57BL/6JJcl mice were purchased from CLEA Japan (Tokyo, Japan) and acclimatized for five days. The mice were exposed to thioacetamide (TAA, TCI, T0817, 300 mg/L) or 4,4'-methylenedianiline (TCI M0220, 750 mg/L) dissolved in drinking water to induce liver damage, whereas the controls received tap water. After four days of treatment, mice were euthanized and the blood was collected through the inferior vena cava into a 1.5 mL tube containing 1.5 µL heparin (Yoshindo). The collected blood sample was centrifuged (1,700 g, 4°C for 15 min) for plasma separation. Plasma alanine aminotransferase (ALT) and aspartate aminotransferase (AST) were measured using a DRI-CHEM NX500sV (Fujifilm).

#### Activation of PSA

Recombinant human pro-PSA (50 µg/mL; R&D Biosystems 1344-SE) was mixed with recombinant thermolysin (1 µg/mL; R&D Biosystems 3097-Zn) in HEPES-Na buffer (100 mM, pH 7.4) containing NaCl (150 mM) and CHAPS (0.1%), and the mixture was incubated at 37°C for 5 min. 1,10-Phenanthroline was added with a final concentration of 20 mM. The activated enzyme solution was either used immediately for the assay or aliquoted for single use, fresh frozen in liquid $N_2$ and maintained at -80°C.

**Activation of granzyme B**

Recombinant human granzyme B (20 µg/mL; R&D Biosystems 2906-SE) was mixed with recombinant mouse cathepsin C (5 mg/mL; R&D Biosystems 2336-CY-) in HEPES-Na buffer (10 mM, pH 5.5), and the mixture was incubated at 37°C for 4 h. The mixture was diluted 1/10 with HEPES-Na buffer (100 mM, pH 7.4) containing NaCl (150 mM) and CHAPS (0.1%). The activated enzyme solution was either used immediately for the assay or aliquoted for single use, fresh frozen in liquid $N_2$ and maintained at -80°C.

**Fluorescence imaging**

Fluorescence images were acquired by microscope (Ti2, Nikon) equipped with a 20× dry objective lens (Plan Apo VC 20×), sCMOS camera (ORCA-Fusion C14440, Hamamatsu Photonics), white LED illumination unit (X-Cite Xylis, Opto Science), and a motorized stage.
The excitation and emission filters used were FITC (mirror = 510 nm, Ex. = 460–500 nm, Em. = 510–560 nm) for Alexa488-conjugated magnetic beads and mCherry (mirror = 600 nm, Ex. = 550–590 nm, Em. = 608–683 nm) for resorufin.

**Preparation of antibody-conjugated beads**

The magnetic beads conjugated with antibodies were prepared using reagents and protocols of the Simoa Homebrew assay starter kit (Quanterix). For fluorescent magnetic beads, the Simoa 488-dyed Singleplex Bead (Quanterix) was used. For PSA, the unconjugated capture antibody of the anti-PSA antibody pair for ELISA (abcam; ab256313) was used. For granzyme B, the unconjugated detector antibody of the anti-granzyme B antibody pair for ELISA (abcam; ab245038) was used. Firstly, the buffer of the antibody solution was exchanged to Bedas conjugation buffer (Quanterix) by three rounds of buffer exchange (loading of the 500 µL solution in an Amicon ultra (50K) and centrifuging at 14,000 rcf × 5 min, 25°C). The concentration of the antibody was measured using a Nanodrop One (Thermo Fisher Scientific), and the solution was diluted to 0.2 mg/mL with Antibody conjugation buffer. $4.2\times10^8$ magnetic beads were washed three times with 300 µL Bead wash buffer (Quanterix) by capturing beads on a magnetic stand (Takara Bio). The beads were dissolved in 291 µL Bead conjugation buffer. A 0.1 mg/mL solution of 1-(3-Dimethylaminopropyl)-3-ethylcarbodiimide (EDC) was freshly prepared and 9 µL was added while vortexing. After shaking at 4°C for 30 min, the supernatant was removed, magnetic beads were quickly washed with Bead conjugation buffer, and 300 µL antibody solution (0.2 mg/mL) was added while vortexing. The mixture was incubated at 4°C for 2 h. The beads were washed with 300 µL Bead wash buffer twice, and shaken in 300 µL Bead blocking buffer (Quanterix) at 25°C for 45 min. The beads were washed with 300 µL Bead wash buffer and stored in 300 µL Bead diluent buffer (Quanterix).

**Fluorophosphonate (FP)-biotin-based enzyme labeling and analysis**

The sample was diluted in HEPES-Na buffer (100 mM, pH 7.4) containing NaCl (150 mM) and CHAPS (0.1%), FP-biotin (Santa Cruz sc-215056) was added with a final concentration of 20 µM, and the reaction was incubated at 25°C for 14 h. Then, the sample was diluted to 1/100 in HEPES-Na buffer (100 mM, pH 7.4) containing NaCl (150 mM) and CHAPS (0.1%), and a 100 µL volume was mixed with 25 µL of antibody-conjugated beads ($2\times10^7$ beads/mL; diluted with Bead diluent buffer (Quanterix)) in a 96-well plate (Thermo Fisher Scientific 249944) and shaken (800 rpm, 30°C) for 30 min using a Microplate shaker (Quanterix). The plate was then washed with a microplate washer (BIOTEK 405 TS). To the remaining beads, 100 µL of streptavidin β-Gal (100 pM, diluted with SBG Diluent buffer; Quanterix) was added and incubated at 30°C for 10 min. The plate was washed with the microplate washer (BIOTEK 405 TS). In SR-X (Quanterix), the beads were mixed with a solution of resorufin β-Gal (10 µM) in HEPES-Na buffer (100 mM, pH 7.4) containing $CaCl_2$ (1 mM), $MgCl_2$ (1 mM) and Triton-X-100 (250 µM) and loaded into a Simoa disk (Quanterix) following the standard protocol. After incubation at 25°C for 1 h, fluorescence images of the Simoa disk were acquired using an epifluorescence microscope.

**FP-azide-based enzyme labeling, click conjugation, and analysis**

FP-azide (ActivX #88316) was added to a sample diluted in HEPES-Na buffer (100 mM, pH 7.4) containing NaCl (150 mM) and CHAPS (0.1%) at a final concentration of 40 µM and the reaction was incubated at 25°C for 18 h. The sample was then diluted 1/10 in HEPES-Na buffer (100 mM, pH 7.4) containing NaCl (150 mM) and CHAPS (0.1%), and alkyne-PEG4-biotin (10 µM), $CuSO_4$ (1 mM), BTTP (3 mM), and sodium ascorbate (1 mM). In the control condition, alkyne-PEG4-biotin (10 µM) was added without other reagents and the reaction was incubated at 25°C for 2 h. The sample was then diluted 1/10 in HEPES-Na buffer (100 mM, pH 7.4) containing NaCl (150 mM) and CHAPS (0.1%), and a 100 µL volume was mixed with 25 µL of antibody-conjugated beads ($2\times10^7$ beads/mL; diluted with Bead diluent buffer (Quanterix)) in a 96-well plate (Thermo Fisher Scientific 249944) and shaken (800 rpm, 30°C) for 30 min using the Microplate shaker (Quanterix). The plate was then washed with the microplate washer (BIOTEK 405 TS). To the remaining beads, 100 µL of streptavidin β-Gal (100 pM, diluted in SBG Diluent buffer; Quanterix) was added, and incubated at 30°C for 10 min. The plate was washed with the microplate washer (BIOTEK 405 TS). In SR-X (Quanterix), the beads were mixed with resorufin β-Gal (10 µM) in HEPES-Na buffer (100 mM, pH 7.4) containing $CaCl_2$ (1 mM), $MgCl_2$ (1 mM) and Triton-X-100 (250 µM) and loaded into a Simoa disk (Quanterix) following the standard protocol. After incubation at 25°C for 1 h, fluorescence images of the Simoa disk were acquired using an epifluorescence microscope.

**Digital ELISA**

Digital ELISA to detect PSA was performed using the Simoa technology (Quanterix) following the manufacturer's protocol. The sample was diluted to 1/100 in HEPES-Na buffer (100 mM, pH 7.4), and 100 µL was mixed with 25 µL of antibody-conjugated beads ($2\times10^7$ beads/mL; diluted with Bead diluent buffer (Quanterix)) in a 96-well plate (Thermo Fisher Scientific 249944) and shaken (800 rpm, 30°C) for 30 min using a Microplate shaker (Quanterix). The plate was then washed with a microplate washer (BIOTEK 405 TS). To the remaining beads, 100 µL of biotinylated secondary-antibody (250 ng/ml) was added and shaken (800 rpm, 30°C) for 10 min using a Microplate shaker (Quanterix). The plate was washed with a microplate washer (BIOTEK 405 TS). To the remaining beads, 100 µL of streptavidin β-Gal (100 pM, diluted in SBG Diluent buffer; Quanterix) was added, and incubated at 30°C for 10 min. The plate was washed with a microplate washer (BIOTEK 405 TS). In SR-X (Quanterix), the beads were mixed with resorufin β-Gal (10 µM) in HEPES-Na buffer (100 mM, pH 7.4) containing $CaCl_2$ (1 mM), $MgCl_2$ (1 mM) and Triton-X-100 (250 µM), and loaded into a Simoa disk (Quanterix) following the standard protocol. After incubation at 25°C for 1 h, fluorescence images of the Simoa disk was acquired using an epifluorescence microscope.

**Enzyme Activity Assay using Microplate Reader**

The enzyme activity assay was performed in HEPES-Na buffer (100 mM, pH 7.4) containing CHAPS (0.1%). The fluorometric assay was performed using half-area 384-well plates (Greiner 784900) (20 µL reaction volume). The absorbance assay was performed using 96-well clear plates (Thermo Fisher Scientific 167008). The signals were acquired with a plate reader, Envision 2103 Multilabel Reader (Perkin Elmer), with appropriate filter settings.

**FP-based enzyme labeling and western blotting**

To the sample diluted in HEPES-Na buffer (100 mM, pH 7.4), FP-biotin was added with a final concentration of 20 µM, and the reaction was incubated at 25°C for 14 h. The samples were diluted in HEPES-Na buffer (100 mM, pH 7.4) and mixed with the same amount of Laemmli sample buffer containing β-mercaptoethanol (2×), and denatured at 95°C for 3 min. For SDS-PAGE, an 8 µL sample (40 µg total protein) was loaded onto a Multigel mini 10/20 (17 wells, Cosmo Bio) and separated using 30 mA/gel constant current condition for 1 h. ECL Rainbow marker (Cytiva) was used as the protein ladder. After electrophoresis, the gels were transferred to PVDF membranes using a semi-dry blotting kit (Protein Transfer Kit for Semidry Electroblot, Cosmo Bio) following the standard protocol. Electrophoresis was performed using 100 mA/gel constant current condition for 1 h. The membrane was washed three times with Tris-buffered saline containing Tween-20 (0.05%; TBS-T), and incubated with Streptavidin-HRP (Cell Signaling Technology #3999, 1/3000 in TBS-T) for 1 h. The membrane was washed three times with TBS-T. The HRP-linked antibody was detected using a chemiluminescence reagent (Westar Supernova, Cyanagen) with a gel imager (ImageQuant LAS 4000 Mini, Cytiva).

**Statistical Analysis**

For data requiring statistical analysis, the experiments were performed with indicated replicates (n), and the data is presented as mean ± S. D. (error bars). Statistical analyses were performed with Student's t-test (two-sided test, equal error variances) using Microsoft Excel.

**Epifluorescence microscopy**

Fluorescence images were acquired using a fluorescence microscope (Ti2, Nikon) equipped with a 20× dry objective lens (Plan Apo 20×), sCMOS camera (ORCA-Fusion C14440, Hamamatsu Photonics), white LED illumination unit (X-Cite Xylis, Opto Science), and a motorized stage. Images were acquired in tile scan mode with perfect focus. The excitation and emission filters used were FITC (mirror = 510 nm, Ex. = 460-500 nm, Em. = 510-560 nm) and mCherry (mirror = 600 nm, ex. = 550-590 nm, Em. = 608-683 nm).

**Image processing**

Images were processed using the GA3 module of NIS Elements software (Nikon). First, all fluorescence images were background-corrected using a rolling ball correction (3 μm). Then, ROIs were chosen by bright spot detection using the FITC and mCherry filters (diameter = 3 μm), and irregular fluorescent spots derived from fluorescent debris or air bubbles were omitted by dilating the ROI and removing the overlapping ROIs by size and shape filter. The number of spots was counted as the number of ROIs in each channel.

### Supplementary figures

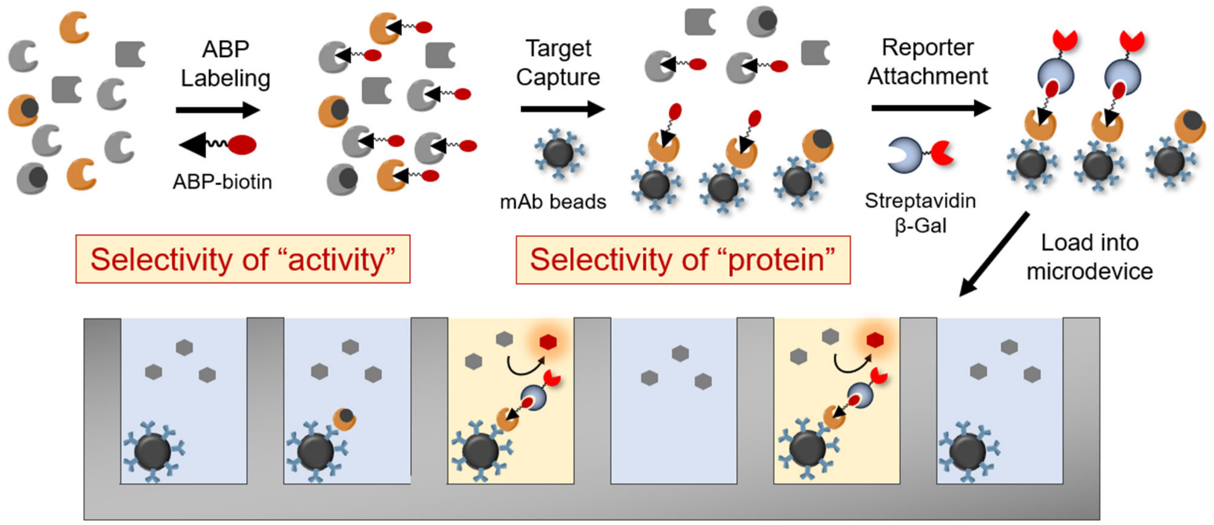


**Figure S1.** Illustrative protocol of single-molecule activity assay to detect active PSA.

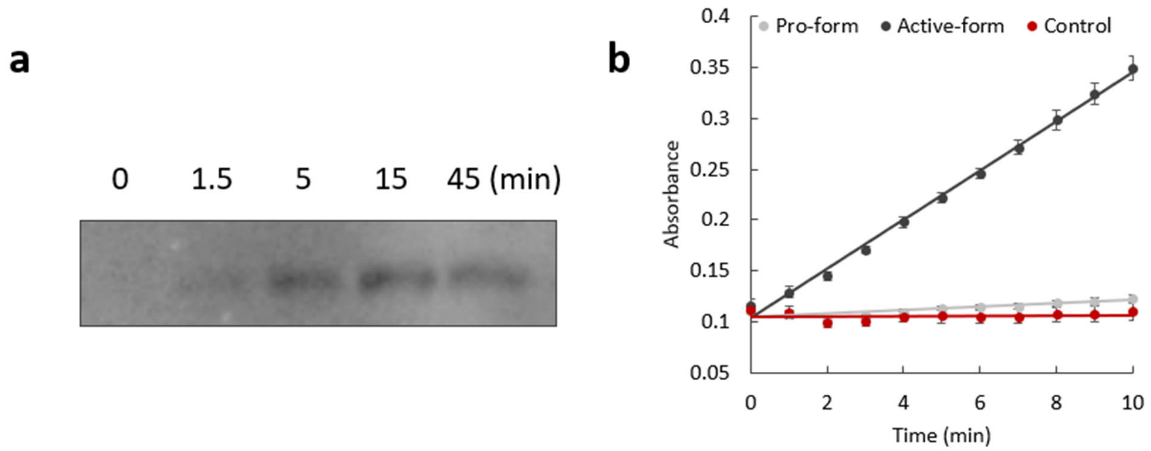


**Figure S2.** Optimization of PSA activation by thermolysin and confirmation of PSA activity using conventional absorption assay. Recombinant human pro-PSA (50 μg/mL) was mixed with recombinant thermolysin (1 μg/mL) in HEPES-Na buffer (100 mM, pH 7.4) containing NaCl (150 mM) and CHAPS (0.1%), and the mixture was incubated at 37°C for 1.5-45 min. 1,10-Phenanthroline was added with the final concentration of 20 mM. The activated enzyme solution was analyzed by Western blotting using Streptavidin-HRP-based chemiluminescence detection. (b) Absorbance (405 nm) change of Suc-RPY-*p*NA (4 mM) after mixing with activated PSA or pro-PSA (10 μg/mL) in HEPES-Na buffer (100 mM, pH 7.4) containing NaCl (150 mM) and CHAPS (0.1%) at 25°C. Error bars represent S. D. (n = 3).

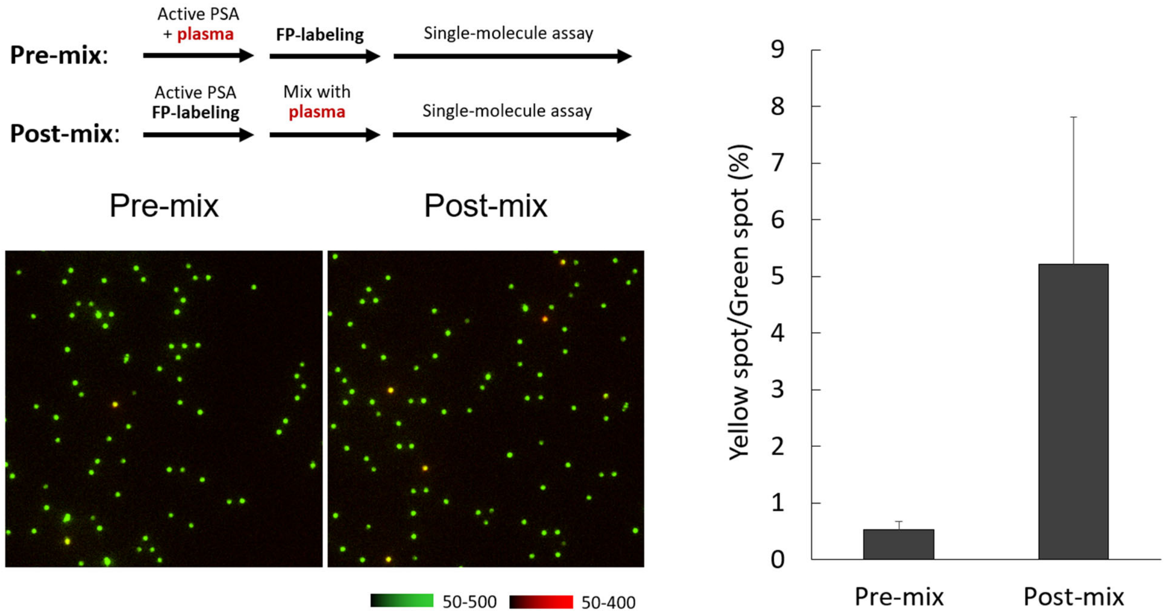


**Figure S3.** Inactivation of PSA activity labeling in plasma samples. For pre-mix conditions, 1/300-diluted plasma samples was mixed with 400 pg/mL activated PSA before labeling with FP-biotin. For post-mix conditions, 400 pg/mL activated PSA was labeled with FP-biotin and mixed with 1/300-diluted plasma samples. The detection method was same as that of F**igure 1b**. Error bars represent S. D. (n = 3).

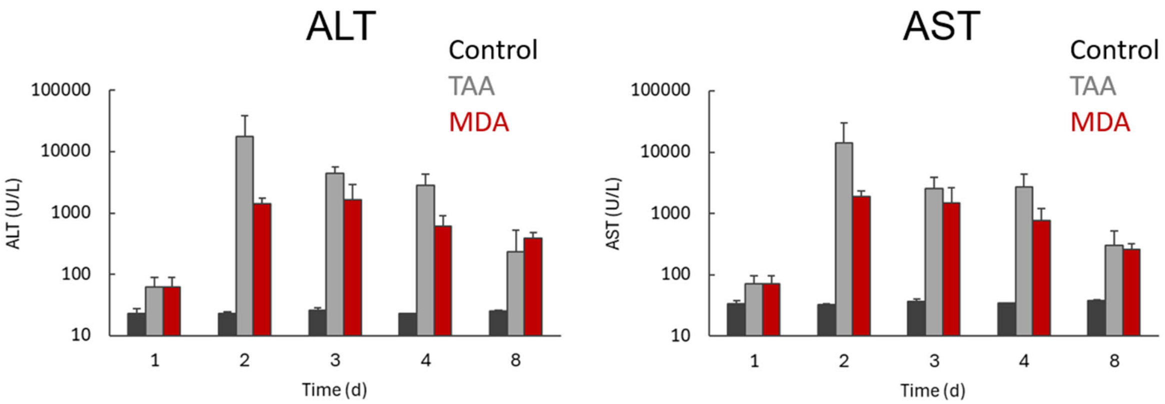


**Figure S4.** AST/ALT of liver damage samples. Error bars represent S. D. (n = 4 for control mice, n = 6 for TAA- and MDA-treated mice).

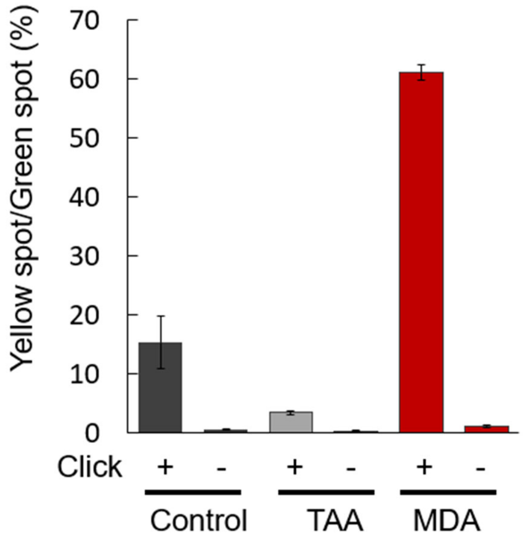


**Figure S5.** Conformation of click-dependent signal in blood samples. The experiment was same with that of **Figure 4d**. Click (-) is the condition in which the click-based elongation of the linker was performed without addition of $CuSO_4$, BTTP, and sodium ascorbate. Error bars represent S. D. (n = 3).
